# Copper impedes calcification of human aortic vascular smooth muscle cells through inhibition of osteogenic transdifferentiation and promotion of extracellular matrix stability

**DOI:** 10.1101/2024.08.30.610428

**Authors:** Iurii Orlov, Gaëlle Lenglet, Carine Avondo, John H. Beattie, Said Kamel, Irina Korichneva

## Abstract

Vascular calcification (VC), a common pathological condition, is a strong predictor of cardiovascular events and associated mortality. Development and progression of VC heavily rely on vascular smooth muscle cells (VSMCs) and are closely related to oxidative stress, inflammation, and remodelling of extracellular matrix (ECM). Copper (Cu), an essential microelement, participates in these processes, however its involvement in pathophysiology of VC and VSMCs physiology remains poorly investigated. In the present study we analysed Cu impact on the calcification of human aortic primary VSMCs induced *in vitro* by treatment with high calcium and phosphate levels. Supplementation with physiological micromolar Cu significantly reduced the amount of calcium deposited on VSMCs as compared to moderate deficiency, Cu restriction with chelators or Cu excess. Moreover, optimal concentrations of Cu ions increased protein production by VSMCs, stimulated their metabolic activity, inhibited alkaline phosphatase activity associated with cell-conditioned medium and cellular lysates, and prevented osteogenic differentiation of VSMCs. RNA-seq results indicated that high calcium and phosphate treatments activated many pathways related to oxidative stress and inflammation in VSMCs at the initial stage of calcification. At the same time, expression of VSMC-specific markers and certain components of ECM were downregulated. Supplementation of calcifying cells with 10 μM Cu prevented most of the transcriptomic alterations induced by high calcium and phosphate while chelation-mediated restriction of Cu greatly aggravated them. In summary, physiological concentration of Cu impedes *in vitro* calcification of VSMCs, prevents their osteogenic transition and minimises early phenotypic alterations induced by high calcium and phosphate, thereby underlining the importance of Cu homeostasis for the physiology of VSMCs, one of the cornerstones of cardiovascular health. Our data suggest that peculiarities of Cu metabolism and its status should be considered when developing preventive and therapeutic approaches for cardiovascular diseases.

## Introduction

Atherosclerosis, defined as a buildup of lipids, cell debris and calcium phosphate in the intimal layer of an arterial wall, leads to the narrowing of large arteries lumen and is the main cause of acute cardiovascular events (Libby et al., 2019). Deposition of calcium phosphate crystals (hydroxyapatite, HA) in the vascular wall, vascular calcification (VC), aggravates atherosclerosis by provoking plaque rupture when localised in the intimal layer or renders arteries stiff and rigid if found in the tunica media (Chow & Rabkin, 2015; Daniela et al., 2020). Irrespective of particular localisation, VC is a strong predictor of cardiovascular events and associated mortality (Rennenberg et al., 2009). VC, which is highly reminiscent of bone formation, is tightly regulated and mainly orchestrated by the predominant cell type in the tunica media, namely vascular smooth muscle cells (VSMCs) (Durham et al., 2018). In normal conditions VSMCs, by means of their coordinated work, control vasodilation and synthesise extracellular matrix (ECM) proteins, generating the flexibility and tensile strength of the vascular wall (Wilson, 2011). However, since VSMCs are not terminally differentiated, they display prominent phenotypic plasticity, and thus can acquire a vast array of phenotypes including pathological ones (Cao et al., 2022). The current paradigm considers VC to be the result of disbalance between calcification inhibitors and activators in favour of the latter. Accordingly, harmful external stimuli may diminish production of molecules inhibiting calcification by VSMCs and provoke them to abundantly secrete pro-calcification factors such as calcium phosphate and different ECM proteins characteristic of bone tissue. VSMCs behaving in such a way concurrently lose specific contractile markers and are deemed to acquire a so-called osteogenic phenotype. This phenotype switch is closely associated with inflammation, oxidative stress and ECM remodelling, often in a positive feedback loop manner (Durham et al., 2018). Better understanding of VC mechanisms, their participants and interactions, may result in novel strategies to prevent deteriorating consequences of VC.

Copper is an essential trace element that controls numerous physiological processes including, but not limited to, cellular respiration and antioxidant defence, maturation of ECM and melanin synthesis (Solomon et al., 2014). Moreover, copper ions directly regulate the activity of central cellular signaling pathways, such as that of the MAPK and PI3K-Akt pathways, and are involved in the control of apoptosis and the cell cycle (Ackerman & Chang, 2018; Grubman & White, 2014; Kardos et al., 2018; Tsvetkov et al., 2022). At the same time unbound copper ions are known to be toxic due to their ability to stimulate ROS formation, provoke protein aggregation and disrupt Fe-S clusters (Liochev & Fridovich, 2002; Macomber & Imlay, 2009; Noda et al., 2013). Coordinated function of specialised copper metabolism system assures proper handling of essential yet potentially toxic copper ions, preventing formation of any free copper (Kaplan & Maryon, 2016). Perturbations of copper metabolism are related to different congenital diseases, development of neurodegenerative disorders and malignancies (Denoyer et al., 2015; Ferreira & Gahl, 2017; Noda et al., 2013).

The role of copper in cardiovascular health remains elusive. On the one hand, higher serum copper is usually reported to be associated with the risk of cardiovascular events (Chen et al., 2015; Huang et al., 2019). However, copper deficiency resulted in myocardial lesions and cardiac hypertrophy (Medeiros, 2017). In its turn, normal dietary intake of copper had an overall beneficial effect on cardiovascular health and mitigated atherosclerosis (Lamb et al., 2001; Wen et al., 2022). The role of copper in the onset and development of VC is not well defined, and so we investigated the impact of physiological copper on the process of *in vitro* calcification in human primary VSMCs and in the modulation of their phenotype.

## Materials and methods

### Extraction and culture of human primary aortic VSMCs

VSMCs were isolated from noncalcified areas of proximal ascending aortas of patients with various cardiovascular diseases at CHU Amiens-Picardie (Amiens, France) by explant method and used from the 3^rd^ to 6^th^ passages. All the procedures were conducted in accordance with the French legislation (ethical agreement number is PI2021_843_0002). Cells were cultured in standard conditions in DMEM (Sigma-Aldrich D6546) supplemented with 15% FBS (Dominique Dutcher Laboratories), Glutamax (Gibco) and 1% penicillin/streptomycin (Sigma-Aldrich). Calcification was induced by 1% FBS DMEM containing 2.2 mM of CaCl_2_ and of phosphate. During experimental procedures cell culture medium was either supplemented by copper in the form of CuCl_2_ or depleted of it by means of copper chelators, namely bathocuproine disulfonate disodium salt (BCS) or ammonium tetrathiomolybdate (TTM). To mimic cuprous form of copper, i.e., Cu(I), AgNO_3_ was utilised. For *in vitro* calcification experiments pro-calcifying medium was changed once every 2 or 3 days, every time it was prepared fresh from cold reagents and equilibrated to ambient temperature before adding to the cells. During calcification biochemical parameters were assessed at timepoints of 7 and 19 days, corresponding to the initial and developed stages of the process, respectively.

### Quantification of deposited calcium

For calcium measurement with o-cresolphthalein complexone, cells were washed with cold PBS and precipitated calcium was dissolved in 0.6 M HCl. The amount of coloured product was measured by absorbance at 565 nm. For the Alizarin red assay, cells were washed and fixed with ethanol, then stained with Alizarin red for 5 minutes. After a quick wash with ethanol, the precipitated dye was dissolved in 10% cetylpiridinium chloride and the absorbance was measured at 560nm.

### MTT assay

After 2 hours of incubation in 1% FBS DMEM containing 0.5 mg/mL MTT, the medium was removed and the cells were air dried for 30 minutes. Formazan crystals were dissolved with DMSO and absorbance read at 570 and 650 nm. The latter value was subtracted from the former, as a mean of background correction.

### Protein extraction and western blot

Proteins were extracted using a buffer based on 1% NP-40 substitute (USBio) containing PMSF and protease inhibitor cocktail (both from Sigma-Aldrich). Scraped cells were shaken at 1000 rpm for 30 minutes at 4 °C. Crude cell lysates were used to determine alkaline phosphatase (ALP) activity while the soluble fraction obtained by centrifugation at 18 000 g for 20 minutes at 4 °C was used for western blotting; protein concentration was measured using DC Protein Assay (Bio-Rad). For western blotting, 25 μg of protein were separated by electrophoresis in 8-20% gradient polyacrylamide gels under denaturing conditions, and transferred onto nitrocellulose membranes which were blocked by 5% skimmed milk in PBST for 30 minutes. Incubation with primary antibodies in 5% skimmed milk in PBST was carried out overnight at 4 °C. After a subsequent 30-minute incubation with secondary HRP-conjugated antibodies in 5% skimmed milk, PBST proteins of interest were detected using the Amersham ECL Select chemiluminescent reagent (Cytiva). The intensity of protein bands was determined using Image Lab software (Bio-Rad). We used a mouse monoclonal anti-ATOX1 antibody (Santa Cruz, sc-100557, 1:1000 dilution), a goat polyclonal anti-SOD1 antibody (Santa Cruz, sc-8637, 1:1000 dilution) and a mouse monoclonal anti-β-actin antibody (Sigma-Aldrich, A1978, 1:5000 dilution) as specific primary antibodies. For detection with secondary antibodies, we used an HRP-conjugated goat anti-mouse antibody (Santa Cruz, sc-2005, 1:5000 dilution) and an HRP-conjugated rabbit anti-goat antibody (Southern Biotech, 6160-05, 1:5000 dilution). Membranes were washed three times for 5-minutes each using PBST between all the described steps.

### ALP activity assay

Samples were mixed with a buffer containing pNPP to achieve the following concentrations of reagents: 10 mM pNPP (Sigma-Aldrich), 1.5 mM MgCl_2_, 50 mM AMP pH 10.5. Absorbance at 405 nm was read immediately after mixing, and after 30-minute incubation at 37 °C. Initial values were subtracted for background correction.

### JC-1 and DHE fluorescent microscopy

Cells grown on glass bottom Petri dishes (Thermo Fisher) were incubated in warm PBS-containing 2 μM JC-1 or 40 μM DHE (Thermo Fisher) for 15 minutes in the dark, then washed with PBS and left to equilibrate in it for 15 more minutes. Fluorescence from live cells was examined at room temperature using a Carl Zeiss confocal microscope system LSM780 (Carl Zeiss Microscopy Gmbh). JC-1 was excited at 488 nm and fluorescence monitored over the following spectral ranges: 516-543 nm (monomeric form) and 579-614 nm (aggregated form). DHE was excited at 488 nm and fluorescence was monitored over the spectral range 550-735 nm. Acquired images were analysed using Zen Blue software v2.6 (Carl Zeiss Microscopy Gmbh).

### RNA extraction, reverse transcription and qPCR

Cellular RNA was extracted using RNeasy kit (Qiagen) following manufacturer’s protocol with an additional step of on-column DNAse digestion. The resulting material was reverse transcribed utilising a High-Capacity cDNA Reverse Transcription Kit (Thermo Fisher Scientific), according to the supplier’s instructions. qPCR was carried out using SYBR green (Eurogentec) as a reporter. Primer sequences are given in **Error! Reference source not found.**.

### Library preparation, RNA sequencing and data analysis

Sequencing of strand-specific libraries was performed on a NextSeq2000 sequencer (Illumina) to obtain 65 bases single-end reads. Analysis of RNA-seq data was performed with Sequana. Briefly, after preprocessing, reads were mapped to the *H. sapiens* reference genome assembly GRCh38 (GCA000001405.28) using STAR. FeatureCounts was used to produce the count matrix using annotation v109 from Ensembl with strand-specificity information. Differential expression testing was conducted using DESeq2 library scripts, indicating the significance (Benjamini-Hochberg adjusted p-values, FDR < 0.05) and the fold-change for each comparison. Enrichment analysis was performed using modules from Sequana. The GO enrichment module utilised PantherDB and QuickGO services. The KEGG pathways enrichment used gseapy, EnrichR and KEGG databases.

### Statistical data analysis and related remarks

Statistical analysis was performed in GraphPad Prism version 8.0.1 (GraphPad Software). Data on figures are represented as mean ± standard deviation, relative to control, which is designated by “CTL” on every figure. Values for deposited calcium, metabolic and ALP activities, are given per unit of protein.

## Results

Copper chloride added to high calcium and phosphate calcification medium (CaCl_2_ and Pi, 2.2 mM of each) resulted in a U-shaped dose-dependent effect on the amount of calcium deposited on VSMCs, indicating that both copper deficiency and its excess result in more profound cellular calcification (Figure 1-A and -B). At the same time, low micromolar concentrations of copper chloride reduced the degree of VSMC calcification, with the concentration of 10 μM being the most efficient. The effect of copper was reversed by both cell-impermeable (BCS) and cell-permeable (TTM) copper chelators (Figure 1-B). Moreover, the observed effect was reproduced by silver ions, which have been shown to some extent to mimic copper ions (Puchkova et al., 2019).

**Figure 1.**
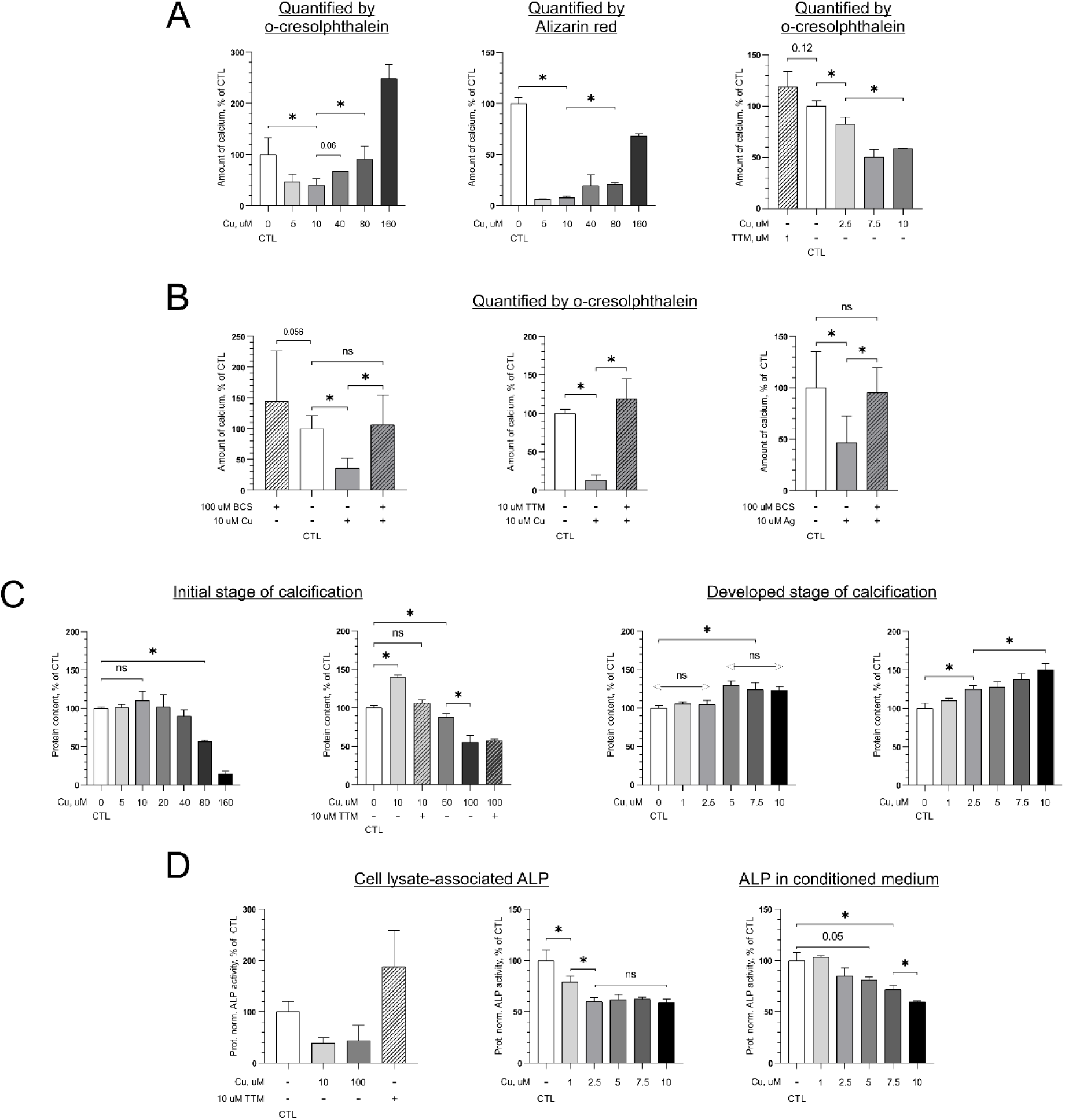
Physiological concentration of copper decreases amount of deposited calcium (A, B), increases cellular protein production (C) and inhibits ALP (D) during calcification of VSMCs induced by high calcium and phosphate (2.2 mM of each). Data are presented as mean ± standard deviation, * designates p-value <0.05 according to the appropriate statistical test. Arrows with “ns” indicate nonsignificant difference according to one-way ANOVA. ALP – alkaline phosphatase; BCS, TTM – copper chelators; CTL – control condition; Cu – copper chloride.

Simultaneously, 10 μM copper resulted in a more active protein synthesis by VSMCs in pro-calcifying medium as compared to moderate restriction of copper (no additional copper in the media), or its substantial excess. This effect increased with time (Figure 1-C). Moreover, we found that 1 μM and above of copper in the pro-calcification medium inhibited ALP activity associated with cell lysates, and copper ≥5 μM had a similar inhibitory effect on ALP in the medium conditioned by VSMCs for 3 days (Figure 1-D).

Figure 2 summarises copper effects on the metabolic phenotype of VSMCs. As such, supplementation with copper resulted in a bell-shaped effect on metabolic activity of VSMCs, according to the MTT assay. Once again, doses of around 10 μM were optimal (Figure 2-A). These changes in metabolic activity at least partially manifested as an increase in the activity of VSMC mitochondria, since incubation of VSMCs with 10 μM copper for 3 days altered the rate of mitochondrial membrane depolarisation, revealed by the JC-1 fluorescent probe (Figure 2-B). Mitochondrial membrane potential, estimated as a ratio of the fluorescence intensities of two forms of JC-1, i.e., red fluorescence of aggregated JC-1 versus green fluorescence of monomeric form, appeared to be higher in the control cells as compared to copper pre-treated cells. Furthermore, under acute oxidative stress, imposed by the addition of 600 μM hydrogen peroxide directly to the cell samples on the microscope stage, increased rate of membrane depolarization of the mitochondria of the cells pre-treated with 10 μM copper was observed, thus supporting our evidence for increased metabolic activity. Such activity is usually accompanied by liberation of reactive oxygen species (ROS). Indeed, our experiments show a tendency towards increased ROS production by VSMC after their incubation with copper. However, VSMCs preincubated with copper demonstrated diminished ROS production in response to acute redox stress mimicked by H_2_O_2_ as estimated by oxidation of redox-sensitive fluorescent probe DHE (**Error! Reference source not found.**). At the same time, low micromolar concentrations of copper significantly increased levels of antioxidant proteins SOD1 and ATOX1 in calcified VSMCs (Figure 2-C). Notably, we did not detect any upregulation of *ATOX1* gene expression at the two examined timepoints (Figure 2-D). Correlation of mitochondrial depolarization with moderate production of ROS would correspond to the phenomenon of pre-conditioning that in turn, by triggering expression of antioxidant proteins, enhances cellular antioxidant defence and stress resistance. Such phenotypic alteration induced by physiological copper may underline its beneficial role for VSMC physiology.

**Figure 2.**
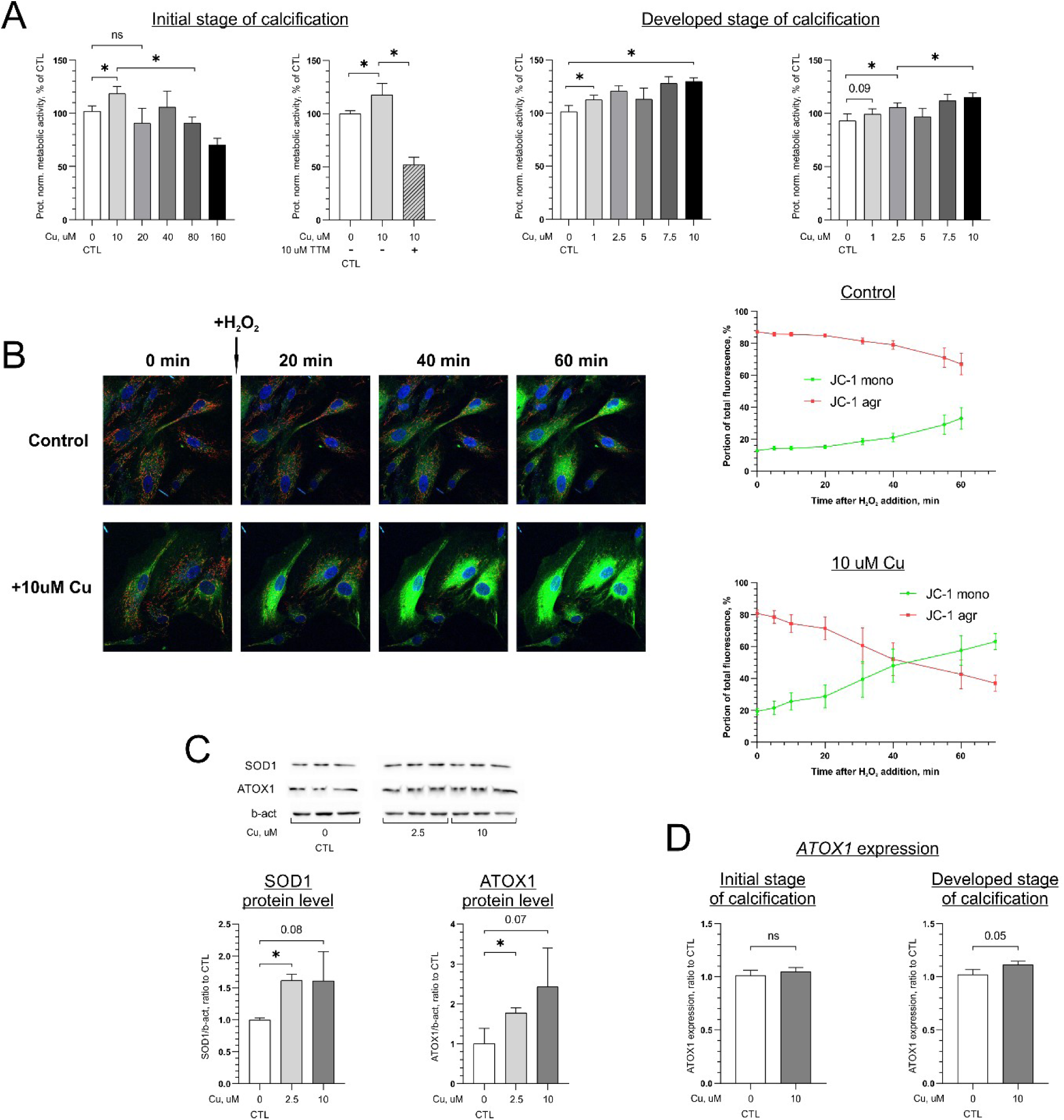
Physiological concentration of copper increases cellular metabolic activity according to MTT assay (A), protein levels of SOD1 and ATOX1 antioxidant proteins (B), though does not influence expression of the latter (D) during calcification of VSMCs induced by high calcium and phosphate (2.2 mM of each). Preincubation with copper renders mitochondria more susceptible to H_2_O_2_-induced depolarisation as revealed by JC-1 life-cell fluorescent microscopy (B). GAPDH was used as a housekeeping gene for qPCR. Data are represented as mean ± standard deviation, * designates p-value <0.05 according to the appropriate statistical test. CTL – control condition; Cu – copper chloride; TTM – copper chelator.

To reveal the potential mechanisms underlying the observed anti-calcification effect of copper, we sequenced transcriptomes of VSMCs calcified in presence of 10 μM copper or 10 μM of the copper chelator TTM, as well as the transcriptomes of the cells incubated in control pro-calcification and no-calcification conditions. Based on the number of differentially expressed genes (DEGs) with substantially altered expression (Log2(|FC|)>1) at the initial stage of calcification (3 days), this process appears to be mainly driven by gene activation and copper reverses these changes (**Error! Reference source not found.**). Specifically, 308 such DEGs (86 and 222 down- and up-regulated, respectively) were detected under calcifying (no added copper) versus non-calcifying condition, whereas 10 μM copper addition to the calcifying medium, reduced nmber of DEGs to 45 (14 and 31 down- and up-regulated, respectively). In contrast, copper depletion by TTM chelation resulted in approximately 1000 DEGs (560 and 405 down- and up-regulated, respectively), indicating significant transcriptomic alterations caused by severe copper deficiency. The 10 most altered DEGs for mentioned conditions are shown in **Error! Reference source not found.**, **Error! Reference source not found.** and **Error! Reference source not found.**, respectively.

According to the gene set enrichment analysis (GSEA), during the initiation of calcification pro-calcifying conditions led to upregulation of pathways related to inflammation and immune stress in VSMCs (Figure 3-A). At the same time, inherent VSMC functions, such as contraction and ECM maintenance, appeared to be supressed (Figure 3-D). Addition of 10 μM copper substantially reversed these effects by inhibiting the upregulation of inflammation-related pathways and preserving a healthy VSMCs phenotype (Figure 3-B and E). We determined that out of 36 gene sets upregulated in control calcification condition, activation of 26 of them was at least partially prevented by copper while a half of gene sets downregulated by pro-calcifying medium were at least partially recovered by copper supplementation (Figure 3-C and F). Copper deprivation by chelation with 10 μM TTM clearly had an opposite effect to copper supplementation. Accordingly, to an even greater extent than high calcium and phosphate alone, TTM induced numerous proinflammatory responses and inhibited physiological VSMC functions (Figure 4-A and C), likely reflecting consequences of severe copper depletion. Thus 11 gene sets from a total of 36 upregulated by high calcium and phosphate alone, were additionally activated by copper deprivation, and 13 out of 15 downregulated sets were even more severely inhibited (Figure 4-B and D). Selected data provided by RNA-seq were validated by qPCR (**Error! Reference source not found.**, **Error! Reference source not found.**). Evidently, copper ions facilitated preservation of contractile marker *ACTA2* and ECM proteins (*COL1A1* and *COL4A1*) gene expression, while they prevented upregulation of the *BMP2* osteogenic differentiation marker.

**Figure 3.**
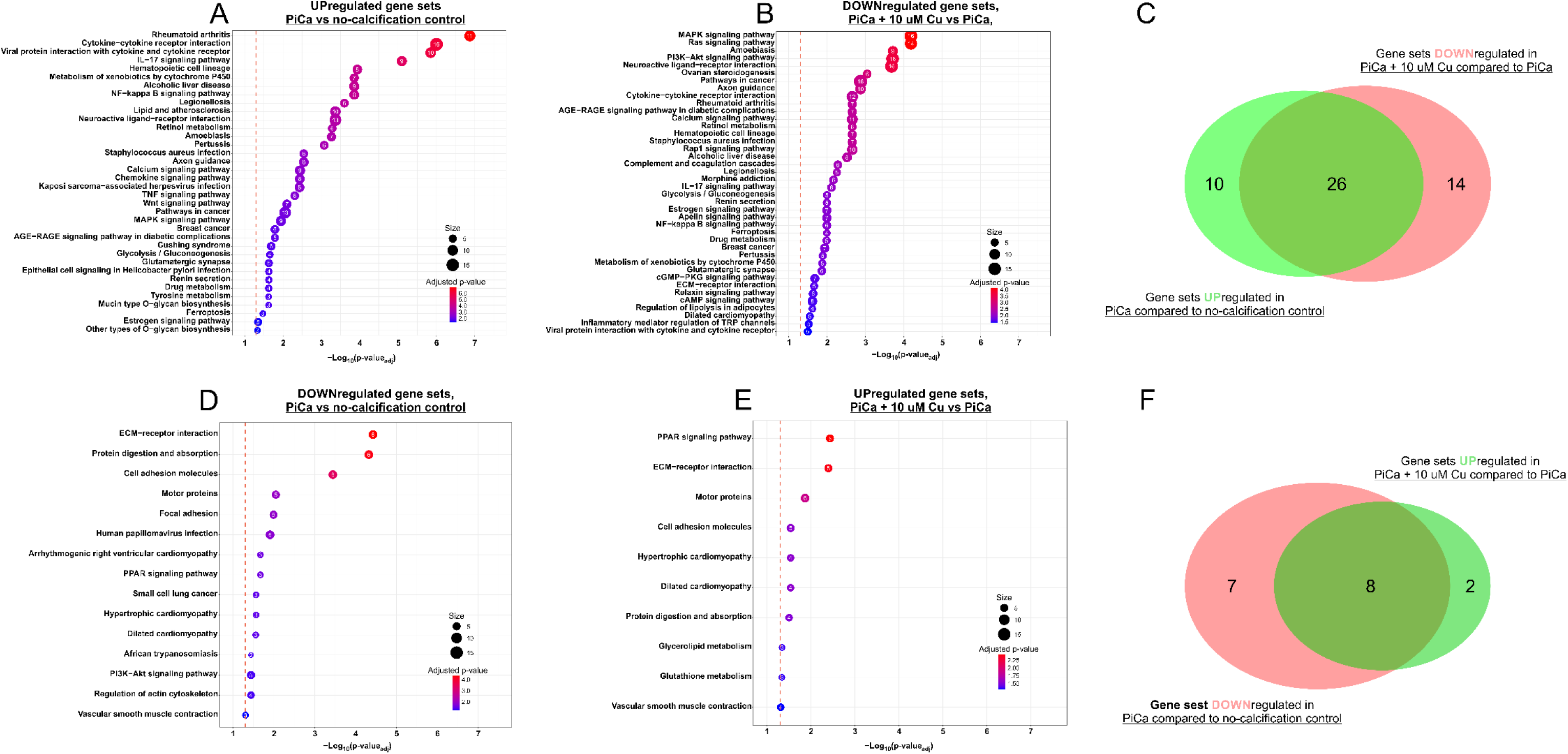
Physiological concentration of copper reverse transcriptomic alterations induced by high calcium and phosphate (2.2 mM of each) at the initial stage (3 days) of VSMCs calcification. <u>A, D</u>: gene sets up- and downregulated, respectively, in VSMCs after their incubation in pro-calcifying medium compared to control condition where no additional calcium or phosphate were added. <u>B, E</u>: gene sets down- and upregulated, respectively, in VSMCs after their incubation in pro-calcifying medium supplemented with 10 uM copper compared to pro-calcifying medium alone. <u>C, F</u>: Venn diagrams representing number of gene sets shared between A-B and D-E, respectively. Cu – copper chloride; PiCa – pro-calcifying medium.

**Figure 4.**
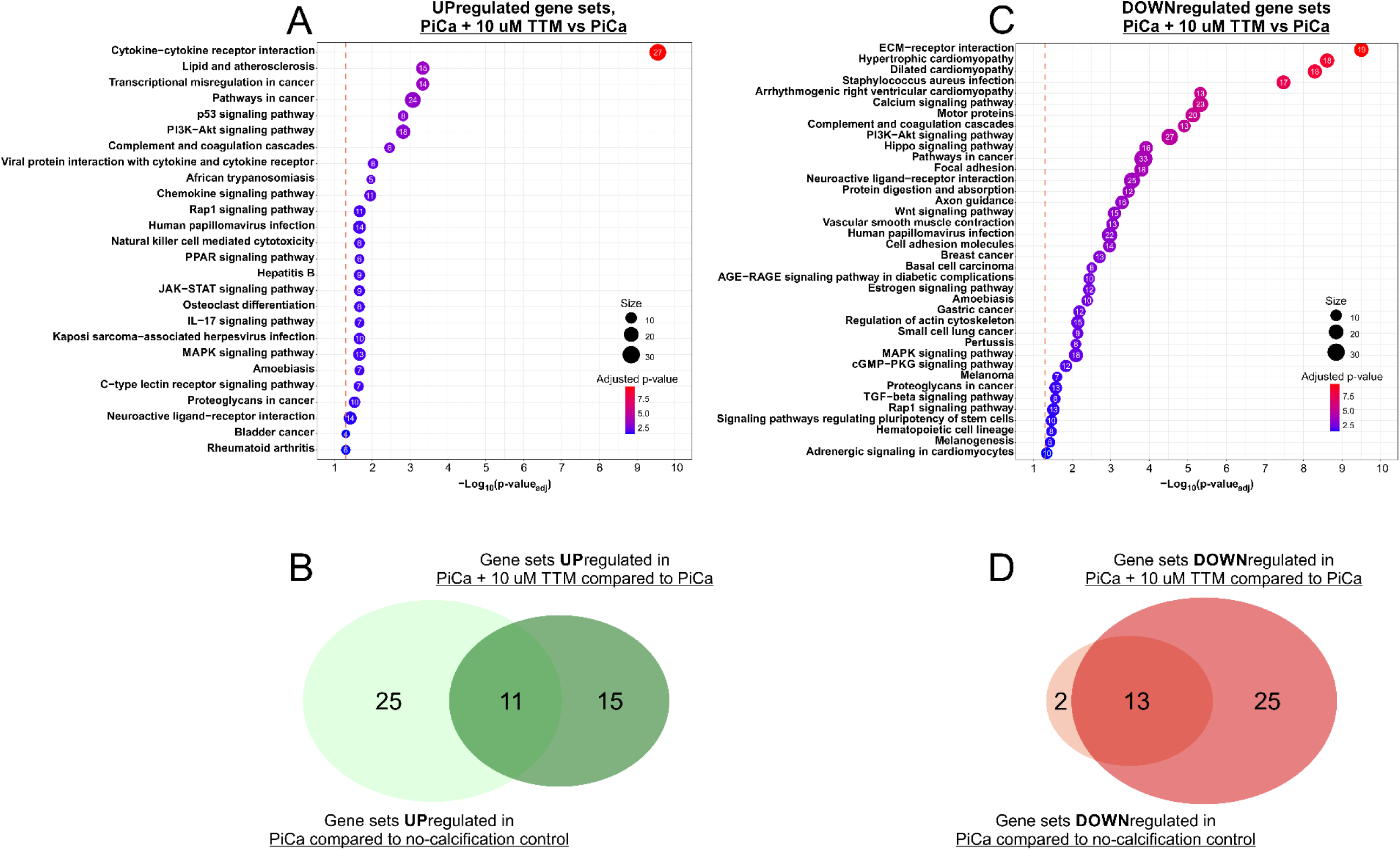
Copper chelation aggravates transcriptomic alterations induced by high calcium and phosphate (2.2 mM of each) at the initial stage of VSMCs calcification. <u>A,</u> <u>C</u>: gene sets up- and downregulated, respectively, in VSMCs after their incubation in pro-calcifying medium with 10uM copper chelator TTM compared to pro-calcifying medium alone. <u>B, D</u>: Venn diagrams representing number of gene sets upregulated by pro-calcifying medium alone that are additionally upregulated in condition with TTM (B) and downregulated by pro-calcifying medium alone that are additionally downregulated in condition with TTM (D). PiCa – pro-calcifying medium; TTM – copper chelator.

**Figure 5.**
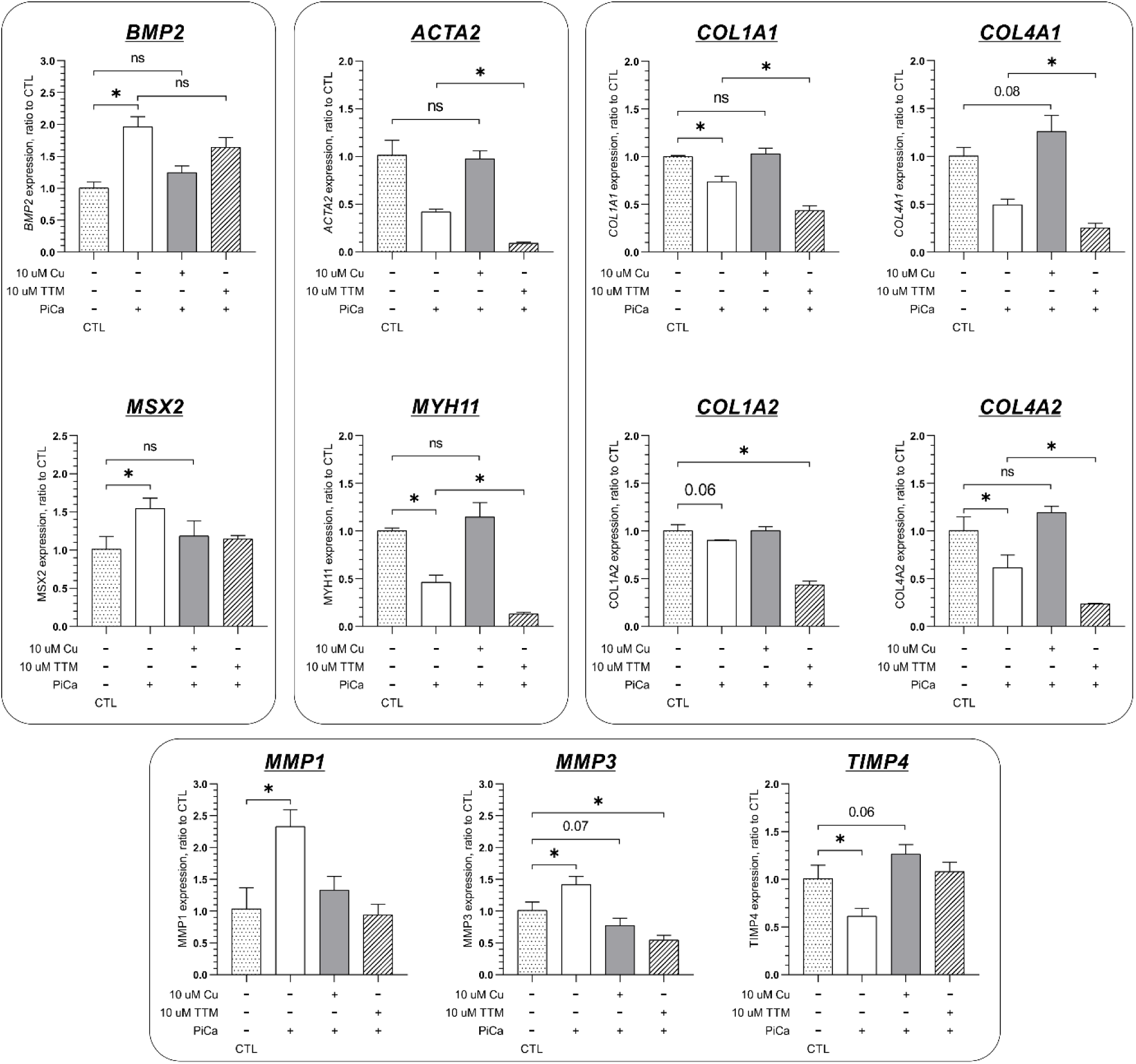
According to qPCR analysis, on gene level physiological concentration of copper prevents upregulation of VSMCs osteogenic transition drivers, preserves expression of VSMCs contractile markers and extracellular matrix proteins at the initial stage of VSMCs calcification induced by high calcium and phosphate (2.2 mM of each). GAPDH was used as a housekeeping gene. Data are represented as mean ± standard deviation, * designates p-value <0.05 according to the appropriate statistical test. CTL – control condition; Cu – copper chloride; PiCa – pro-calcifying medium; TTM – copper chelator.

The results were different when calcification was already well developed (9 days). When using 1%FBS DMEM condition as reference, higher numbers of DEGs (Log2(|FC|)>1) in copper supplemented cells (400 and 264 down- and up-regulated, respectively) compared to what was found for the cells incubated in control calcifying medium with no copper added (89 and 124 down- and up-regulated, respectively) (**Error! Reference source not found.**). However, GSEA did not indicate any downregulated gene sets in cells calcified in presence of 10 μM copper, compared to control cells calcifying without copper supplementation. Number of upregulated genes sets were limited to 11 with the majority related to carbohydrates and lipids metabolism (**Error! Reference source not found.**). At the same time copper chelation by 10 uM TTM at the developed stage of calcification led to the greatest among all our experiments number of DEGs (**Error! Reference source not found.**). Overall, severe copper deprivation resulted in a marked dysregulation of gene expression with a multitude of disturbed pathways (**Error! Reference source not found.**).

## Discussion

The results of our study show that physiological concentrations of copper in the pro-calcifying medium reduces the amount of deposited calcium during calcification of VSMCs and changes their phenotype by increasing protein synthesis and overall metabolic activity. Physiological copper was found to facilitate preservation of VSMCs contractile markers, inhibit their osteogenic differentiation, promote ECM stability, and prevent upregulation of inflammation and immune response-related pathways by environment favouring calcification. At present, the reports on copper involvement in the calcification process are limited. It has been demonstrated that copper ions can integrate into HA crystals and thus facilitate their dissolution, extracts of copper-doped HA powders showed lower pro-calcification activity, as compared to un-doped, according to representative images of rat mesenchymal stem cells stained with Alizarin Red, and copper addition decreased osteogenic differentiation-related RUNX2 expression in MC3T3-E1 upon their growth on copper-doped hydroxyapatite bioceramics (Bazin et al., 2021; Noori et al., 2024). Taking present scarcity of data, our work brings new knowledge in the largely understudied field and, to our best knowledge, is the first account of copper significance for VC.

It is important to note that in our study cells were supplemented with Cu(II). Although the main cellular copper gateway, CTR1, is Cu(I)-specific, and thus requires reduction of Cu(II) prior its import, evidence exists that Cu(II) can be directly imported through other transporters, most notably DMT1 (Lin et al., 2015). Copper speciation inside the cell is still an area of active research and the roles of intracellular copper ions with different oxidation states remain elusive. In our work silver ions, which partly mimic Cu(I), reproduced well anti-calcifying effect of copper (Puchkova et al., 2019). Consequently, cuprous copper can be suggested as the form mainly responsible for the effects observed. Undoubtedly, further studies are required to make a more definitive conclusion.

Increased metabolic activity and protein synthesis by physiological copper during calcification of VSMCs demonstrated in this study may be linked to both enhanced proliferation and/or cellular production of proteins, including structural components of the ECM. In agreement with the latter, our RNA-seq and qPCR data confirm stimulation of ECM gene expression by 10 μM copper. Some of the data pointing to increased metabolic activity may be explained by the effect of copper on mitochondria. Pre-incubation with copper appears to have a pre-conditioning effect on VSMC mitochondria, making them more prone to depolarisation, which in turn may result in higher energy production (**Error! Reference source not found.**). Our supplemental data showing time-courses of ROS production after pre-treatment with copper as compared to the control, reveals a pre-conditioning effect with higher ROS production but slowed redox response after peroxide stimulation of the cells pre-incubated with copper. This hypothesis is corroborated by our data on the induction of antioxidant proteins by copper supplementation.

Healthy, non-disturbed VSMCs abide in a quiescent state characterised by low proliferation and expression of contraction-related markers (Durham et al., 2018). In response to various harmful stimuli, VSMCs can acquire different phenotypes, including those promoting development of pathological processes like calcification (Tyson et al., 2020). Conventionally, two cellular states, contractile and synthetic, were described with the latter demonstrating higher rates of proliferation, migration and protein synthesis as compared to the former. The synthetic state was also considered to foster vascular pathological alterations. However, the existence of a repertoire of cellular phenotypes has recently been suggested (Cao et al., 2022; Yap et al., 2021). While to a varying degree they share some common features, like increased proliferation or protein production, each of them bears their own markers and executes specialised functions. Accordingly, osteogenic-like VSMCs lose contractile markers and abundantly secrete bone-specific components of ECM. This phenotype is known to facilitate VC, and BMP2 is one of the well-established drivers of osteogenic transition (Li et al., 2008; M. Sun et al., 2017). Recently, together with mesenchymal-, macrophage- and adipocyte-like states, researchers identified fibroblast-like VSMCs. They actively express mediators of cell adhesion, ECM organisation as well as proliferation and are primarily responsible for the deposition of collagen-rich ECM (Sorokin et al., 2020; Yap et al., 2021). We reported that copper supplementation 1) inhibits ALP activity 2) prevents induction of pathways mediating immune/inflammation responses and 3) osteogenic differentiation, 4) preserves VSMCs contractile markers and 5) increases expression of genes encoding adhesion-related molecules and collagens. Therefore, we may suggest that in our model, copper supplementation steers VSMCs towards a fibroblast-like phenotype instead of an osteogenic phenotype, or at least impedes the process of osteogenic transition (Kim et al., 2020). Despite association of fibroblast-like VSMCs with certain pathologies, there are reports that increased protein secretion by these cells and enhanced proliferation may facilitate fibrous cap stability during atherogenesis (Cao et al., 2022; Wirka et al., 2019; Yap et al., 2021). Adding that copper was found to stimulate VSMCs migration (Ashino et al., 2018) and our unpublished data) it is reasonable to conclude that physiologically-relevant concentrations of copper may be beneficial in mitigating the progression of atherosclerosis.

The role of copper in cardiovascular health is still controversial. On the one hand, elevated serum copper has been associated with advanced calcification of aortic valve and cornea in patients on maintenance hemodialysis, increased incidence of cardiovascular events, severity of cardiovascular diseases (CVDs) and cardiovascular mortality (Bagheri et al., 2015; Chen et al., 2015; Ford, 2000; Huang et al., 2019; W. Sun et al., 2015). However, higher copper intake was reported to decrease abdominal aortic calcification, positively correlate with the level of circulating cardioprotective protein Klotho and protect against stroke and myocardial infarction (Liu et al., 2023; Ostojic et al., 2023; Wen et al., 2022; Yang et al., 2022). Moreover, in cholesterol-fed rabbits adequate copper intake resulted in smaller atherosclerotic lesions while targeted delivery of copper to already formed lesions reduced their size. (Lamb et al., 2001; Wang et al., 2021). Finally, there is a point of view that copper deficiency is one of the main overlooked causes of CVDs (Klevay, 2016). This controversy may be associated with nuances of different methodologies used to assess copper status. For example, while total copper in circulation heavily depends on ceruloplasmin concentration, other factors such as loosely bound or exchangeable copper as well as activity of blood cell or plasma cuproenzymes may be of great importance by themselves (Hambidge, 2003; Maares et al., 2023).

The reference range for copper in serum of a healthy individual is 10-20 μM. According to different estimates 40-90% of copper in human plasma is bound to ceruloplasmin while the rest is associated with albumin, α2-macroglobulin, SOD3, other proteins and small molecules (Linder, 2016). Within the total plasma copper, several potential copper pools are distinguished depending on the tightness of ion binding, i.e., unexchangeable, loosely bound, exchangeable or labile copper. The amount of plasma copper accessible for cellular uptake is also not well determined, with various estimates in the low micromolar range being reported (Maares et al., 2023). Our data clearly demonstrate that 10 μM copper had the most beneficial effect on VSMCs during their *in vitro* calcification and it likely corresponds to the most physiologically accessible copper in the circulation. Nevertheless, it should be noted that VSMCs are not in direct contact with plasma, however, at early stages of atherosclerosis compromised endothelial permeability may be an important factor. The details of the molecular mechanisms of copper transport to VSMCs remain largely unknown and are subject for future investigation. We believe that our study is a significant step in understanding the importance of copper homeostasis for human VSMC biology and for cardiovascular health in general, assisting the development of personalized approaches to the prevention of cardiovascular diseases.

## Acknowledgements

Library preparation, sequencing, differential gene expression analysis and gene set enrichment analysis were performed by G. Haustant, L. Lemée, E. Kornobis at Biomics Platform, C2RT, Institut Pasteur, Paris, France, supported by France Génomique (ANR-10-INBS-09) and IBISA.

## Conflict of Interest Statement

Authors declare no conflict of interest

## Data Availability Statement

Raw data generated in the current study is available for download from SRA: https://www.ncbi.nlm.nih.gov/sra/PRJNA1039967.

## Statements

This project received funding from the European Union’s Horizon 2020 research and innovation programme under grant agreement No. 734931 (IK) and was supported by Vernadski PhD scholarship from the French government (IO).

### Ethics approval statement

All the procedures were conducted in accordance with the French legislation, ethical agreement number is PI2021_843_0002.

### Patient consent statement

Before the procedure all patients gave a written informed consent for the use of their cellular material for research purposes.

## Notes

### Competing Interest Statement

The authors have declared no competing interest.

