## Supplementary material for "Copper impedes calcification of human aortic vascular smooth muscle cells through inhibition of osteogenic transdifferentiation and promotion of extracellular matrix stability"

### Supplementary data

| Gene | Primer | Sequence, 5'-3' |
|---|---|---|
| **ACTA2** | F | GTGTTGCCCCTGAAGAGCAT |
| | R | GCTGGGACATTGAAAGTCTCA |
| **ATOX1** | F | CTGTGGAGGCTGTGCTGAAG |
| | R | TCTTGTTGGGCAGGTCAATG |
| **BMP2** | F | CGGACTGCGGTCTCCTAA |
| | R | GGAAGCAGCAACGCTAGAAG |
| **COL1A1** | F | GATTCCCTGGACCTAAAGGTGC |
| | R | AGCCTCTCCATCTTTGCCAGCA |
| **COL4A1** | F | GCCCCCCAGGCAGAGA |
| | R | CCAACTCCTTTTCCATCATACTGA |
| **GAPDH** | F | TCGGAGTCAACGGATTTGG |
| | R | GCAACAATATCCACTTTACCAGAGTTAA |

***Table S1. Sequence of primers used for qPCR analysis.***

| | | Any $Log_2(\|FC\|)$ | | $Log_2(\|FC\|)>1$ | |
|---|---|---|---|---|---|
| **Condition 1** | **Ref. cond.** | **Down** | **Up** | **Down** | **Up** |
| **PiCa** | **noPiCa** | 2627 | 2457 | 86 | 222 |
| **PiCa+10Cu** | **noPiCa** | 1251 | 1372 | 14 | 31 |
| **PiCa+10TTM** | **noPiCa** | 4170 | 4404 | 560 | 405 |
| **PiCa+10Cu** | **PiCa** | 3075 | 3211 | 285 | 119 |
| **PiCa+10TTM** | **PiCa** | 4238 | 4208 | 678 | 417 |

***Table S2. Number of differentially expressed genes between various experimental conditions at the initial stage of VSMCs calcification (3 days). Cu – copper chloride; $Log_2(|FC|)$ – base 2 logarithm of an absolute value of expression fold change between condition 1 and reference condition; noPiCa – 1% FBS DMEM; PiCa – pro-calcifying medium, i.e. 1% FBS DMEM, 2.2 mM of each $CaCl_2$ and Pi; TTM – ammonium tetrathiomolybdate; VSMCs – vascular smooth muscle cells.***

| $Log_2(FC)$ | Gene name | Description |
|---|---|---|
| **5.7** | CYP1A1 | cytochrome P450 family 1 subfamily A member 1 |
| **5.0** | VIPR1 | vasoactive intestinal peptide receptor 1 |
| **4.4** | NKD2 | NKD inhibitor of WNT signaling pathway 2 |
| **4.4** | KRT32 | keratin 32 |
| **4.3** | SEMA5B | semaphorin 5B |
| **4.0** | ALOX15 | arachidonate 15-lipoxygenase |
| 4.0 | CYP1B1 | cytochrome P450 family 1 subfamily B member 1 |
| **3.8** | SECTM1 | secreted and transmembrane 1 |

| | | |
|---|---|---|
| **3.7** | RIPOR3 | RIPOR family member 3 |
| **3.6** | SHISA2 | shisa family member 2 |

***Table S3. Top 10 genes with mostly altered expression between control calcification and no-calcification conditions at the early stage of calcification. Genes which activation is reversed by copper addition are underscored. Calcification was induced by 1% FBS DMEM, containing 2.2 mM of each $CaCl_2$ and Pi. 1% FBS DMEM was used as no-calcification control. $Log_2(FC)$ – base 2 logarithm of expression fold change.***

| $Log_2(FC)$ | Gene name | Description |
|---|---|---|
| **5.2** | AKR1B10 | aldo-keto reductase family 1 member B10 |
| **4.4** | AKR1B15 | aldo-keto reductase family 1 member B15 |
| **2.8** | NQO1 | NAD(P)H quinone dehydrogenase 1 |
| **-2.6** | ADRA2B | adrenoceptor alpha 2B |
| **-2.4** | ANGPTL4 | angiopoietin like 4 |
| **2.4** | CACNA1G | calcium voltage-gated channel subunit alpha1 G |
| **2.2** | OSGIN1 | oxidative stress induced growth inhibitor 1 |
| **2.2** | SMIM43 | small integral membrane protein 43 |
| **2.2** | MT1G | metallothionein 1G |
| **2.1** | SPTA1 | spectrin alpha, erythrocytic 1 |

***Table S4. Top 10 genes with mostly altered expression between conditions of calcification in presence of 10 uM copper and no-calcification control at the early stage of calcification. Calcification was induced by 1% FBS DMEM, containing 2.2 mM of each $CaCl_2$ and Pi. 1% FBS DMEM was used as no-calcification control. $Log_2(FC)$ – base 2 logarithm of expression fold change; TTM – ammonium tetrathiomolybdate.***

| $Log_2(FC)$ | Gene name | Description |
|---|---|---|
| **-7.4** | MYH1 | myosin heavy chain 1 |
| **-5.8** | LEP | leptin |
| **-5.4** | TNNT2 | troponin T2, cardiac type |
| **-5.1** | MYH2 | myosin heavy chain 2 |
| **5.0** | CEACAM1 | CEA cell adhesion molecule 1 |
| **4.6** | DRAXIN | dorsal inhibitory axon guidance protein |
| **-4.4** | LY6G6C | lymphocyte antigen 6 family member G6C |
| **-4.4** | COL11A1 | collagen type XI alpha 1 chain |
| **-4.4** | MYH11 | myosin heavy chain 11 |
| **-4.2** | RSPO1 | R-spondin 1 |

***Table S5. Top 10 genes with mostly altered expression between conditions of calcification in presence of 10 uM copper chelator TTM and no-calcification control at the early stage of calcification. Calcification was induced by 1% FBS DMEM, containing 2.2 mM of each $CaCl_2$ and Pi. 1% FBS DMEM was used as no-calcification control. $Log_2(FC)$ – base 2 logarithm of expression fold change.***

| Gene | Experiment | PiCa | 10 uM Cu | 10 uM TTM |
|---|---|---|---|---|
| **BMP2** | **RNA-seq, mean** | 1.71 | 0.99 | 1.98 |

| | | | | |
|---|---|---|---|---|
| | **qPCR, mean±SD** | 1.96±0.16 | 1.25±0.11 | 1.64±0.15 |
| **MSX2** | **RNA-seq, mean** | 1.43 | 1.01 | 1.67 |
| | **qPCR, mean±SD** | 1.55±0.13 | 1.18±0.20 | 1.15±0.05 |
| **ACTA2** | **RNA-seq, mean** | 0.44 | 0.97 | 0.13 |
| | **qPCR, mean±SD** | 0.42±0.03 | 0.98±0.08 | 0.09±0.01 |
| **MYH11** | **RNA-seq, mean** | 0.34 | 0.98 | 0.05 |
| | **qPCR, mean±SD** | 0.46±0.08 | 1.15±0.15 | 0.13±0.02 |
| **COL1A1** | **RNA-seq, mean** | 0.67 | 0.99 | 0.65 |
| | **qPCR, mean±SD** | 0.74±0.06 | 1.03±0.06 | 0.44±0.05 |
| **COL1A2** | **RNA-seq, mean** | 0.78 | 1.00 | 0.64 |
| | **qPCR, mean±SD** | 0.88±0.12 | 1.02±0.06 | 0.43±0.05 |
| **COL4A1** | **RNA-seq, mean** | 0.47 | 1.13 | 0.37 |
| | **qPCR, mean±SD** | 0.49±0.06 | 1.26±0.17 | 0.25±0.05 |
| **COL4A2** | **RNA-seq, mean** | 0.58 | 1.07 | 0.35 |
| | **qPCR, mean±SD** | 0.62±0.13 | 1.19±0.07 | 0.24±0.00 |
| **MMP1** | **RNA-seq, mean** | 4.19 | 1.06 | 1.78 |
| | **qPCR, mean±SD** | 2.33±0.26 | 1.33±0.22 | 0.94±0.17 |
| **MMP3** | **RNA-seq, mean** | 1.08 | 1.02 | 3.95 |
| | **qPCR, mean±SD** | 1.42±0.13 | 0.78±0.11 | 0.55±0.08 |
| **TIMP4** | **RNA-seq, mean** | 0.25 | 1.02 | 1.50 |
| | **qPCR, mean±SD** | 0.6±0.15 | 1.28±0.08 | 1.05±0.05 |

***Table S6. Comparison of expression level values obtained by RNA-seq and qPCR for some genes between different experimental conditions at the initial stage of calcification. $Log_2(FC)$ is indicated, 1% FBS condition is taken as a reference level in all the cases. GAPDH was used as a housekeeping gene for qPCR. Calcification was induced by 1% FBS DMEM, containing 2.2 mM of each $CaCl_2$ and Pi. Cu – copper chloride; $Log_2(FC)$ – base 2 logarithm of expression fold change; PiCa – pro-calcifying medium; TTM – ammonium tetrathiomolybdate.***

| | | **Any $Log_2(\lvert FC\rvert)$** | | **$Log_2(\lvert FC\rvert)>1$** | |
|---|---|---|---|---|---|
| **Condition 1** | **Ref. cond.** | **Down** | **Up** | **Down** | **Up** |
| **PiCa** | **noPiCa** | 2545 | 2912 | 89 | 124 |
| **PiCa+10Cu** | **noPiCa** | 3622 | 4025 | 400 | 264 |
| **PiCa+10TTM** | **noPiCa** | 5926 | 6082 | 2531 | 2172 |
| **PiCa+10Cu** | **PiCa** | 2332 | 2548 | 53 | 38 |
| **PiCa+10TTM** | **PiCa** | 5857 | 5929 | 2282 | 1947 |

***Table S7. Number of differentially expressed genes between various experimental conditions at the developed stage of VSMCs calcification (9 days). Cu – copper chloride; $Log_2(\lvert FC\rvert)$ – base 2 logarithm of an absolute value of expression fold change between condition 1 and reference condition; noPiCa – 1% FBS DMEM; PiCa – pro-calcifying medium, i.e. 1% FBS DMEM, 2.2 mM of each $CaCl_2$ and Pi; TTM – ammonium tetrathiomolybdate; VSMCs – vascular smooth muscle cells.***

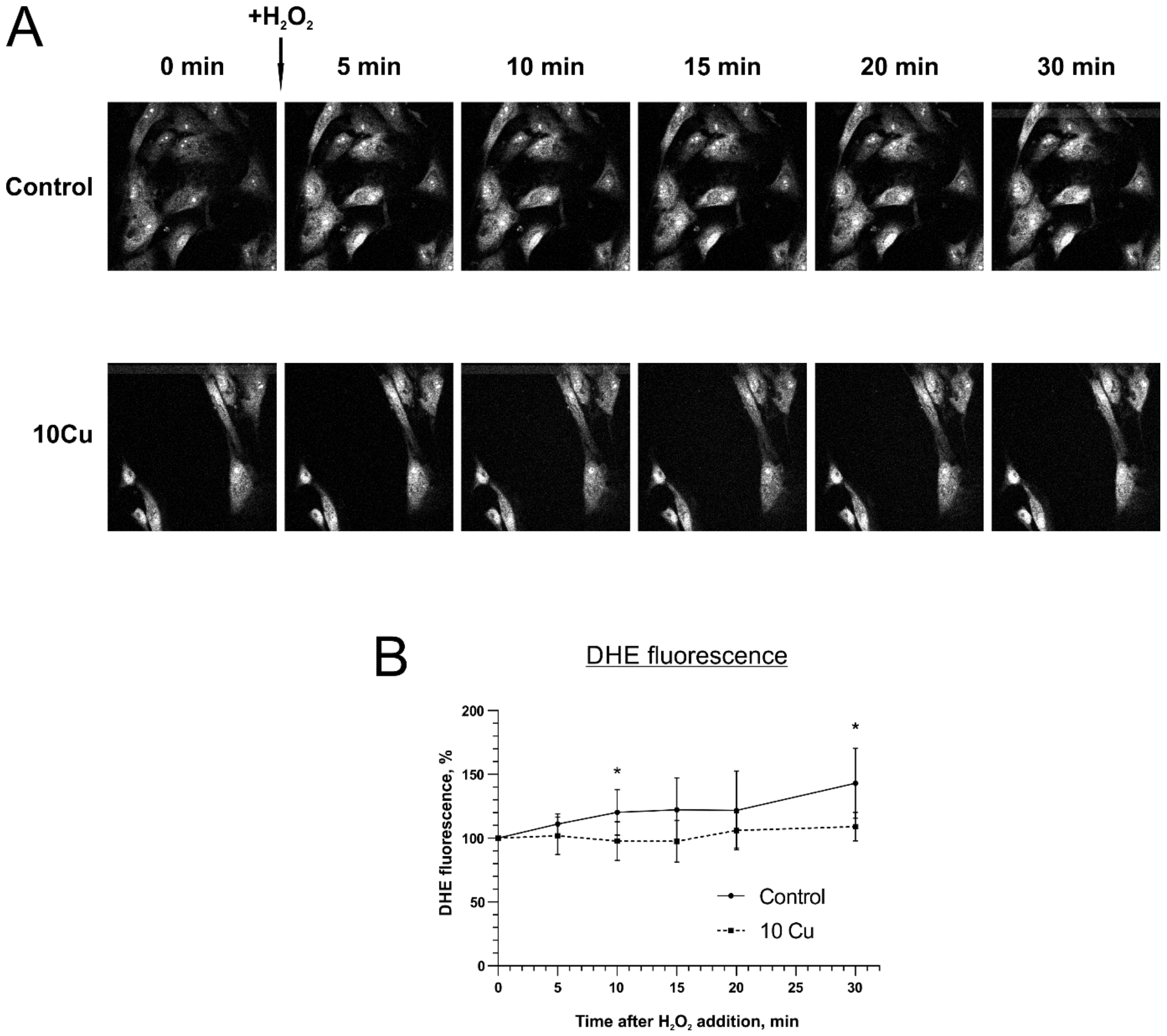


***Figure S1. Preincubation of VSMCs with 10 uM copper leads to diminished intracellular ROS production induced by acute redox stress (1 mM $H_2O_2$) as estimated by oxidation of redox-sensitive fluorescent probe DHE. Fluorescent images of VSMCs loaded with DHE (A) and quantification of its fluorescence (B). Data are represented as means ± standard deviation, * designates p-value <0.05 according to the appropriate statistical test; DHE – dihydroethidium.***

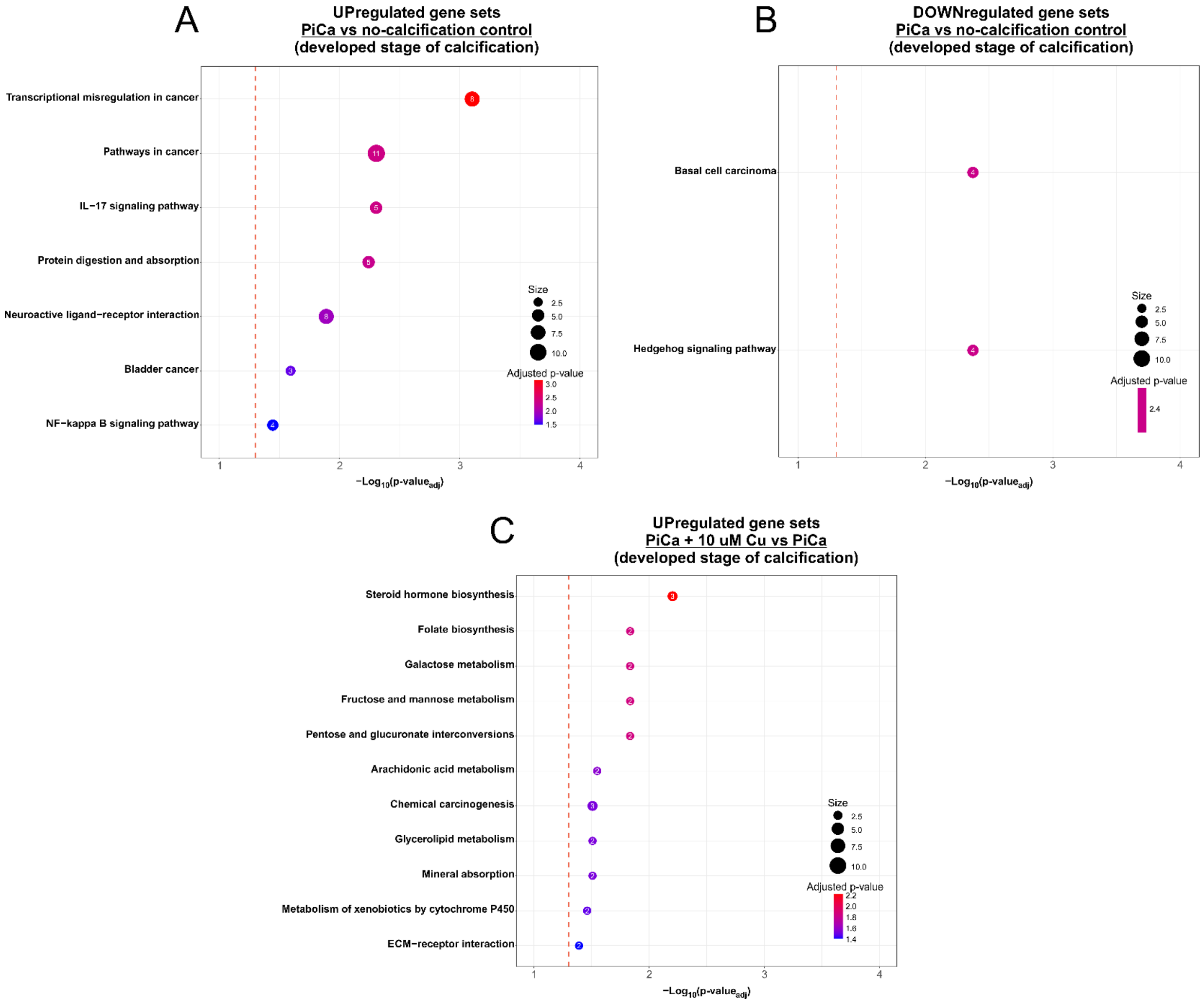


***Figure S2. Supplementation with physiological concentration of copper cannot completely prevent transcriptomic alterations induced by high calcium and phosphate (2.2 mM of each) as shown by gene set enrichment analysis at the developed stage of VSMCs calcification. A, B: gene sets up- and downregulated, respectively, in VSMCs after their incubation in pro-calcifying medium compared to control condition where no additional calcium or phosphate were added. C: gene sets upregulated in VSMCs after their incubation in pro-calcifying medium supplemented with 10 uM copper compared to pro-calcifying medium alone. There were no downregulated genes for this comparison. Cu – copper in the form of copper chloride; PiCa – pro-calcifying medium.***

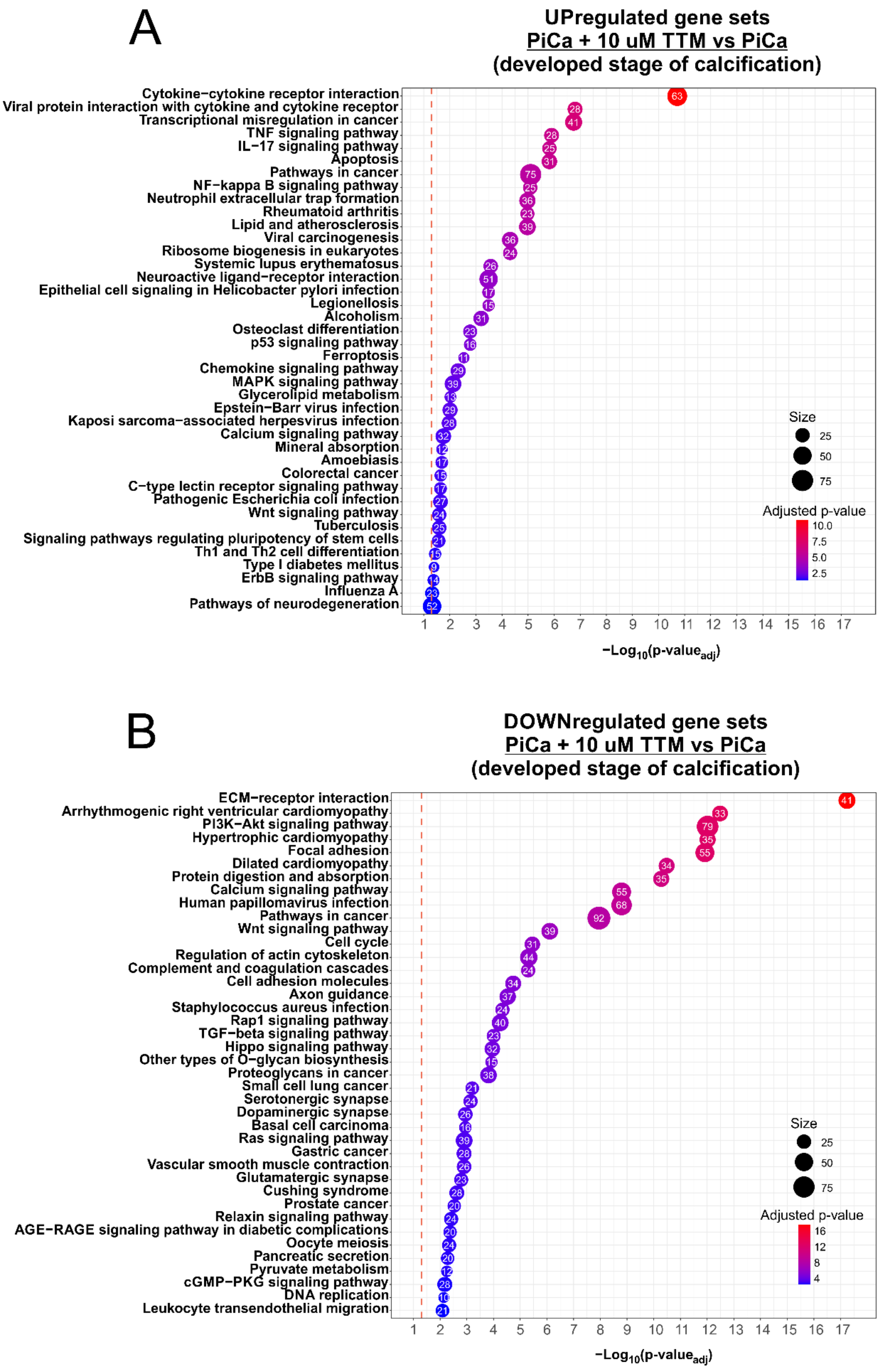


*Figure S3. Copper chelation by 10 uM TTM severely aggravates transcriptomic alterations induced by high calcium and phosphate (2.2 mM of each) as shown by gene set enrichment analysis at the developed stage of VSMCs calcification. A, B: gene sets up- and downregulated, respectively, in VSMCs after their incubation in pro-calcifying medium with 10 uM TTM compared to control calcification condition. PiCa – pro-calcifying medium; TTM – ammonium tetrathiomolybdate.*
